# Impact of temporal pH fluctuations on the coexistence of nasal bacteria

**DOI:** 10.1101/2020.09.15.298778

**Authors:** Sandra Dedrick, M. Javad Akbari, Samantha Dyckman, Nannan Zhao, Yang-Yu Liu, Babak Momeni

## Abstract

To manipulate nasal microbiota for respiratory health, we need to better understand how this microbial community is assembled and maintained. Previous work has demonstrated that the pH in the nasal passage experiences temporal fluctuations. Yet, the impact of such pH fluctuations on nasal microbiota is not fully understood. Here, we examine how temporal fluctuations in pH might affect the coexistence of nasal bacteria. We take advantage of the cultivability of nasal bacteria to experimentally assess their responses to pH. Based on experimentally observed responses, we formulate a mathematical model to numerically investigate the impact of temporal pH fluctuations on species coexistence. Through extensive numerical simulations, we find that the composition of nasal communities is robust against pH fluctuations. Our results suggest that nasal microbiota could be more robust than expected against environmental fluctuations.

## Introduction

Resident microbes in the human nasal passage protect us from respiratory pathogens [1,2]. Indeed, previous research shows the role of resident commensals in suppressing pathogens, such as *Staphylococcus aureus* [3–5]. Investigating how this microbial community is formed and maintained can therefore provide powerful insights into microbiota-based therapies to prevent or treat infections. While such an investigation appears formidable in complex environments such as the gut microbiota, it is feasible for nasal microbiota. First, the nasal microbiota has relatively low diversity, with the majority of composition often attributed to 3~8 species [6]. Second, the majority of these species are readily culturable aerobically *in vitro* under controlled environments [6,7]. Third, both the species and the nasal environment can be sampled relatively easily [8,9].

Many factors, including interspecies interactions [1], the host immune system [10], and resource availability and access [11] can impact the nasal microbiota. However, all these factors take place in an environment that may fluctuate over time and vary in space. Previous investigations have revealed that the nasal passage is in fact very heterogeneous, both spatially and temporally [9]. In particular, pH fluctuations (in the range of 5.8-7.2, depending on the sampling site and time) were observed within the nasal passage [12,13]. Previous studies also demonstrated that temporal environmental fluctuations can increase and support biodiversity based on the temporal niche partitioning mechanism; i.e., environment variations creates additional niches and allow for more species to coexist [14,15]. The purpose of our work is not to introduce a new theoretical framework for modeling microbial communities. Instead, we aim for a predictive mathematical model to study the impact of temporal pH fluctuations on the nasal microbiota composition. Other factors notwithstanding, we specifically ask whether, and when, incorporating temporal pH fluctuations is necessary to accurately predict compositional outcomes. To answer this question, we first use *in vitro* communities constructed from nasal isolates to quantify the community response to temporal pH variations. Then, with parameters relevant to nasal microbiota, we use a phenomenological model to represent microbes and their interactions in an environment with a temporally fluctuating pH. Based on our empirical characterizations of nasal bacteria, we construct *in silico* examples of nasal microbiota and quantify their response to temporal pH fluctuations. Our simulation results suggest that temporal pH fluctuations do not have a major impact on the stable coexistence of nasal bacteria. The outline of our procedure to assess the impact of temporal pH fluctuations on nasal microbiota is shown in Fig 1.

**Fig 1.**
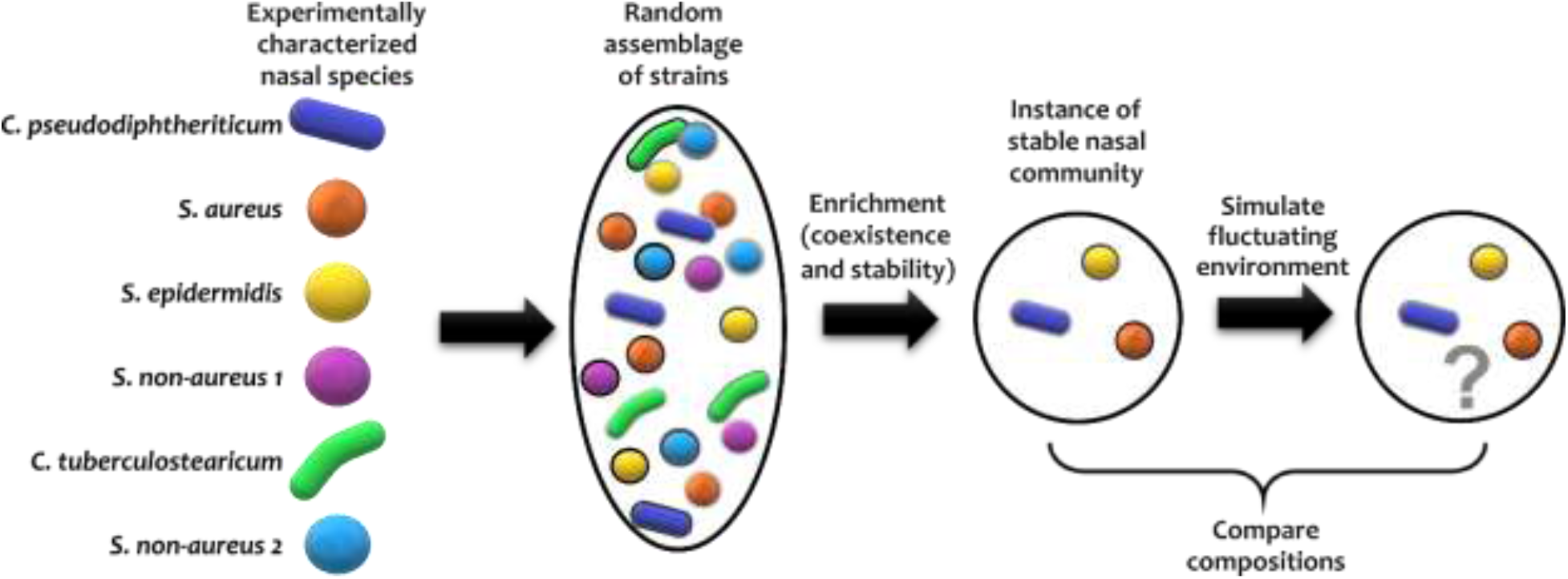
The outline of the procedure for assessing the impact of temporal pH fluctuations on nasal microbiota is shown. To assemble *in silico* nasal communities, we characterized 6 bacterial nasal isolates. We then created *in silico* strains by randomly modulating the parameters of each characterized strain— schematically illustrated as different shades for each species. Using random assemblies of such strains, we simulated the enrichment process to find instances of stable nasal communities. We exposed these communities to a fluctuating pH and compared how the community composition was affected.

## Results

### *In vitro* characterization of nasal bacteria

We experimentally characterized how six representative nasal bacterial strains respond to different pH values in their environment. These bacterial strains were chosen from a set of isolates (see Methods) based on three major considerations: (1) they reliably grow in our cultivation media under an aerobic environment; (2) they include commonly observed *Staphylococcus* and *Corynebacterium* species; and (3) they span the phylogenetic landscape of both closely and distantly related bacteria found in the nasal environment [6]. We assumed that each of these characterized strains is a representative strain of the corresponding species.

We first characterized the pH response of each strain by growing them under different environmental pH values. Different strains exhibited different degrees of pH dependency in their growth rates and carrying capacities (Fig 2). Among these strains, *S. epidermidis*, *S. non-aureus 1*, and *S. non-aureus 2* show fairly similar growth properties. We chose to treat these as separate species in our investigation, because—as shown later—they had considerably different interactions with other species (Fig 3).

**Fig 2.**
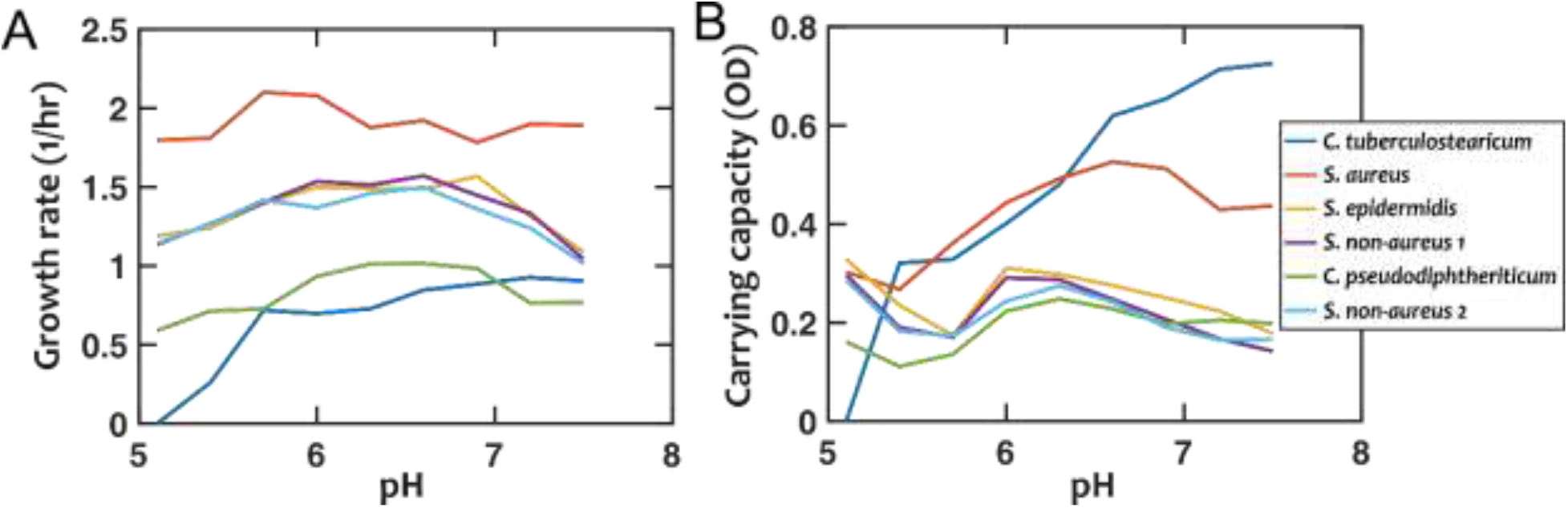
Growth properties of nasal bacterial isolates are pH-dependent. Growth is characterized using the growth rate in the early exponential phase (A), and the carrying capacity based on optical density (OD, absorption measured at 600 nm) as a proxy (B). Each data point is the average of at least 6 replicates from two independent experiments. Error-bars are not shown to avoid overcrowding the plot but the values are available in the raw data. In all cases, growth is experimentally tested in a 10-fold diluted Todd-Hewitt broth with yeast extract (10% THY).

We then examined how different species interacted with one another. For this, we grew each species to its stationary phase in a monoculture, filtered out the cells, and measured how other strains grew in the resulting cell-free filtrates (see Methods). From these measurements, we estimated the inter-species interaction coefficients based on the generalized Lotka-Volterra model (Fig 3; see Methods). In this formulation, baseline competition with complete niche overlap will result in an interaction coefficient of −1. Of note, from our experimental data we cannot distinguish the relative contribution of competitive niche overlap and interspecies facilitation. Nevertheless, for simplicity we only use “facilitation” for extreme cases in which facilitation outweighs competition and the interaction coefficient turns positive. We interpret different gradations of negative interaction coefficients from −1 to 0 as different degrees of niche overlap (with −1 indicating complete niche overlap), and cases with interaction coefficients less than −1 indicate inhibition beyond competition for resources. Among the 30 pairwise interaction coefficients, there were 3 positive values (bright blue, marked by ‘+’ in Fig 3). For simplicity, throughout this manuscript, we assume that these interaction coefficients are not pH-dependent.

**Fig 3.**
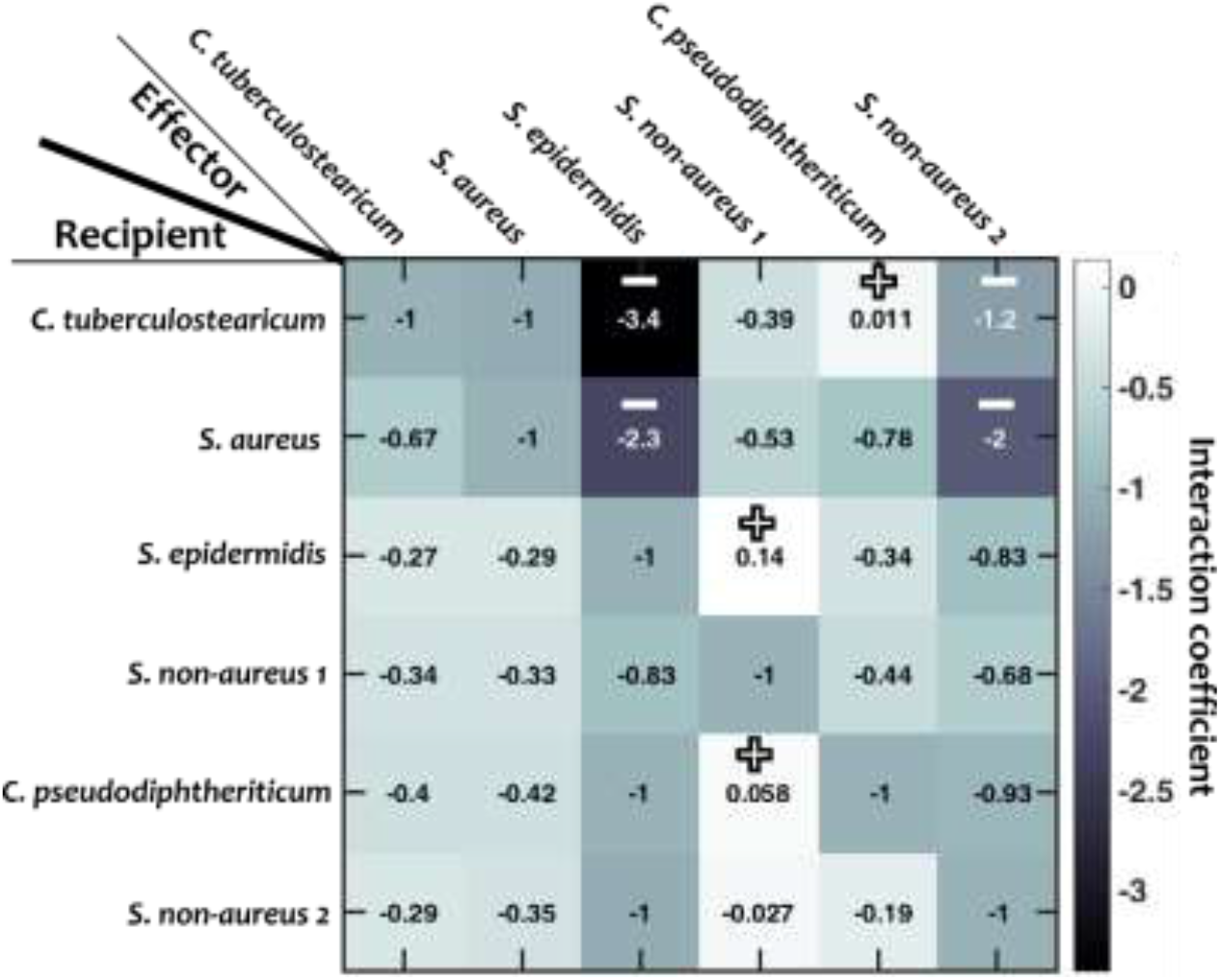
Interaction coefficients among pairs of nasal bacteria. Values represent interaction coefficients in a Lotka-Volterra model. In each case, the growth of a recipient strain is measured when the strain is exposed to cell-free filtrate derived from the effector strain. Positive coefficients (indicating facilitation) and negative coefficients below −1.2 (indicating strong inhibition) are marked by ‘+’ and ‘−’ respectively.

### *In silico* assembly of nasal bacterial communities

To capture some of the diversity of nasal microbiota, we propose that other *in silico* strains of each species can be constructed by randomly modulating the measured properties of that species (i.e. growth rate, carrying capacity, and interaction coefficients). We chose the degree of strain-level modulation to be up to 20%, as a balance between intraspecies and interspecies diversity (Fig S1).

To assess the response of nasal microbiota to temporal fluctuations in the environment, we first construct an ensemble of *in silico* communities that represent a subset of possible nasal communities. This is chosen as an alternative to performing an *in vivo* study, because performing these experiments with human subjects is not feasible and there is no reliable animal model for human nasal bacteria. Compared to *in vitro* studies, these *in silico* communities give us full control over confounding factors and allows us to examine the mechanisms contributing to sensitivity to pH fluctuations [16]. To construct *in silico* communities, we mimicked enrichment experiments [17,18] by simulating the dynamics of an initial assemblage of 20 strains (sampled from the space of *in silico* strains) until the community reached stable coexistence. These *in silico* communities were largely robust against experimental noise in characterization (Fig S2). The interspecies interactions in our model appear to be instrumental in the assembly of these *in silico* communities, as evidenced by changes when we assigned the interaction coefficients at given levels (Fig S3A) or modulated the measured interactions (Fig S3B). To assess how pH fluctuations in the environment influence nasal communities, we take several instances of *in silico* nasal communities, expose them to a fluctuating pH, and quantify how the community composition is affected. The entire process is outlined in Fig 1.

### *In silico* nasal communities are diverse and favor facilitation

We first examined the properties of assembled *in silico* communities at various pH values with no temporal fluctuations. We found that the prevalence of different species were distinct and pH-dependent (Fig S4). This prevalence is a result of nasal species’ pH-dependent growth properties as well as their interspecies interactions.

We also found that during the process of assembling *in silico* communities, the prevalence of interspecies facilitation interactions increased. Comparing the prevalence of facilitation in initial assemblages of strains versus the final stable communities, we found that among the communities that had at least one facilitation interaction at the start of the *in silico* enrichment (89% of communities), facilitation was enriched in ~66% of the final community assemblies (Fig S5).

### Temporal pH fluctuations only minimally impact nasal microbiota composition

Next, we asked how the temporal variation in the environment might influence the community composition. To answer this question, we used instances of *in silico* communities to evaluate the impact of temporal pH fluctuations. We assumed a continuous growth situation in which all community members experience a constant dilution rate. This dilution mimics the turnover in microbiota, for example, when the mucosal layer gets washed away. To avoid situations in which the *in silico* community itself was not stable, we changed the dilution rate by ±50% and only kept the communities for which the modified dilutions only caused a small deviation in community composition (see Methods). Indeed, we found that communities with compositions more sensitive to dilution rates are also more sensitive to pH fluctuations (Fig S6). In all cases, composition deviations were calculated using the Bray-Curtis dissimilarity measure (see Methods).

To evaluate the impact of pH fluctuations, we simulated a controlled sinusoidal pH variation over time, with two parameters: the amplitude and frequency of temporal variations. Thus, *p*(*t*) = *p*_0_ + ΔpH sin(2*πf_pH_t*). Keeping the frequency of fluctuations fixed *f*_pH_ = 0.2/hr), we observed that the deviation in population composition increased with an increasing pH fluctuation amplitude (ΔpH). However, the resulting dissimilarity in population composition was mostly minor, with >85% of cases showing less than 0.2 dissimilarity even when the amplitude of pH fluctuation was set to 1 (Fig 4A). We then examined the impact of the frequency of pH variations, while we kept the amplitude of pH fluctuations fixed (ΔpH = 0.5). At intermediate frequencies, the pH fluctuations caused the largest dissimilarity in community composition compared to stable communities with fixed pH (Fig 4B).

**Fig 4.**
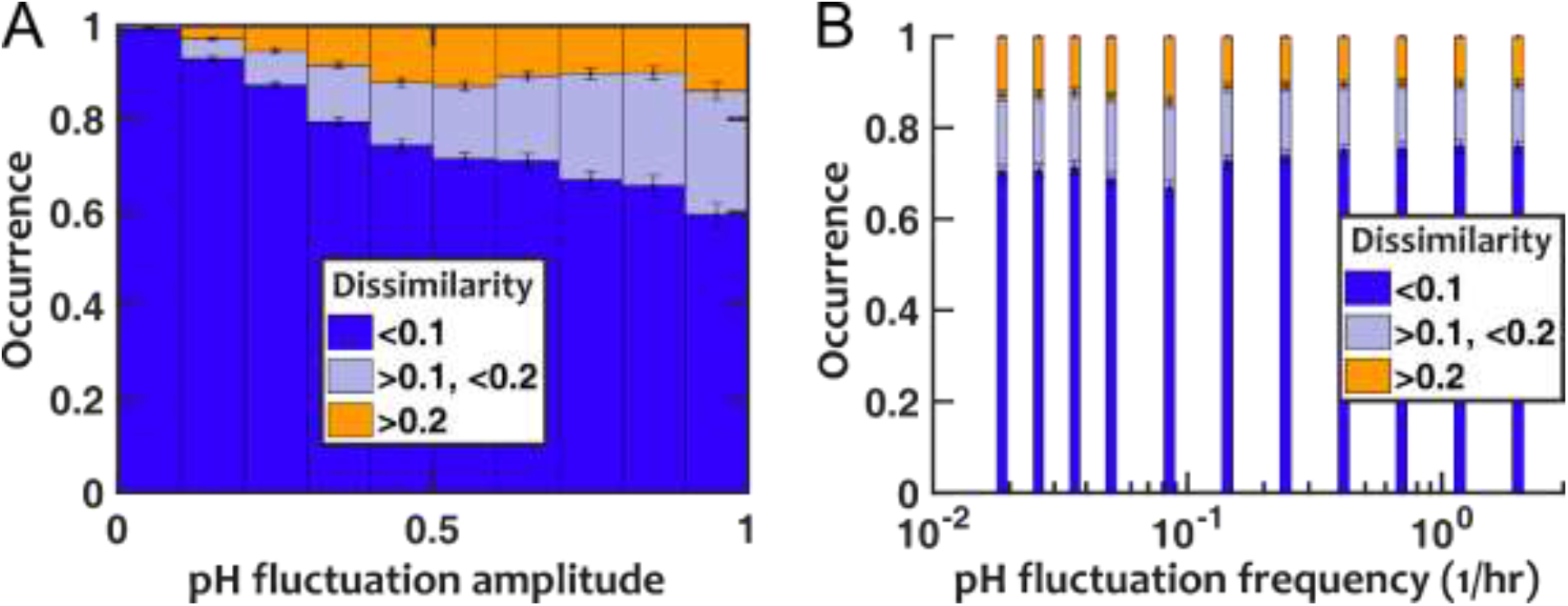
Nasal microbiota composition is robust against pH fluctuations. (A) For a fixed fluctuation frequency (*f*_pH_ = 0.2/hr), larger fluctuation amplitudes increase how the community composition deviates from the no-fluctuation steady state (as quantified with composition dissimilarity). (B) The impact of temporal fluctuations is maximum at intermediate frequencies. Here, the pH fluctuation amplitude is fixed (ΔpH = 0.5). Number of *in silico* communities examined for each condition: n = 10,000.

We repeated the assessment of pH fluctuations by assuming a pH that randomly fluctuated between two discrete pH values to ensure that our results were not limited to sinusoidal fluctuations. The results were overall consistent with sinusoidal pH fluctuations (Fig S7): (1) larger pH fluctuation amplitudes increased the deviation in population composition, but overall the majority of communities only experienced modest deviations; and (2) pH fluctuations at intermediate frequencies had the largest impact on community composition.

### Interspecies facilitations dampen the impact of temporal fluctuations

To explain the low sensitivity of community composition to pH fluctuations, we hypothesized that interspecies facilitation stabilizes the composition by creating interdependencies within the community. From the data in Fig 4, we picked and compared communities with low (“competitive,” 0% facilitation) and high (“cooperative,” 50% facilitation) prevalence of facilitation. The results show that cooperative communities have a consistently and significantly lower composition deviation when exposed to temporal pH fluctuations (Fig 5).

**Fig 5.**
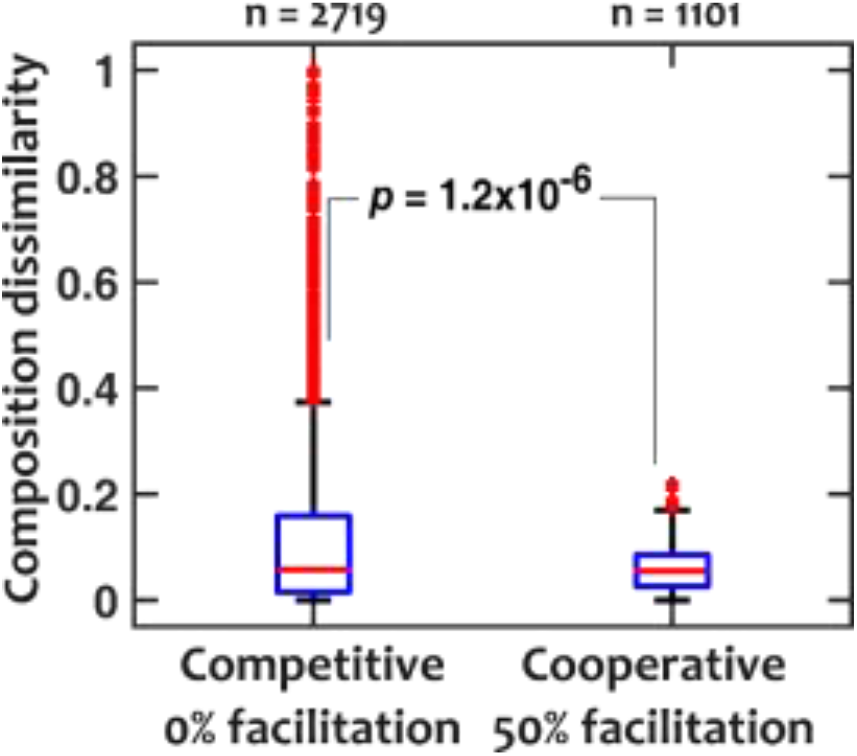
Cooperative communities are more robust against pH fluctuations compared to competitive communities. For competitive (those with 0% facilitation among members) and cooperative (those with 50% facilitation interactions among members) communities, dissimilarity medians were 0.057 and 0.055 and dissimilarity means were 0.12 and 0.058, respectively (*p* = 1.2×10^−6^ with a Mann-Whitney U test). pH fluctuates sinusoidally with a frequency of *f_pH_* = 0.2/hr and an amplitude of ΔpH = 0.5.

To further explore the impact of facilitation, we asked how interspecies niche overlap (the magnitude of negative interspecies interactions) and prevalence of facilitation (the fraction of interspecies interactions that are positive) contribute to sensitivity to pH fluctuations. In our results, we found that larger interspecies niche overlap leads to more sensitivity to pH fluctuations (Fig 6A). This trend holds except when interspecies niche overlap approaches 1; at such high overlaps the community loses diversity (Fig 6B), becoming less sensitive to pH fluctuations. When we directly changed the prevalence of facilitation, we observed that with higher prevalence of facilitation the communities became more diverse and less sensitive to pH fluctuations (Figs 6C and 6D).

**Fig 6.**
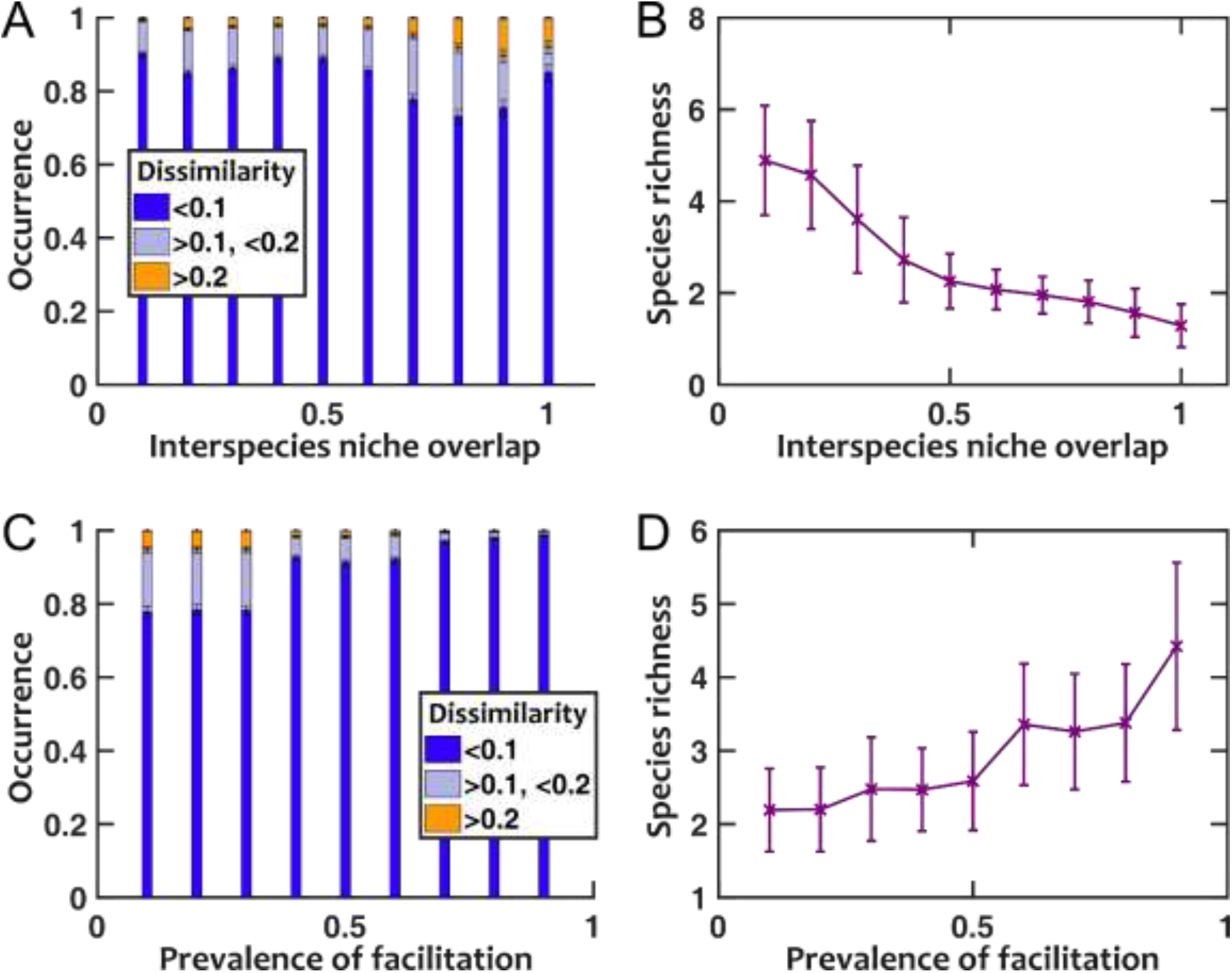
Lower niche overlap and more prevalent facilitation decrease the sensitivity to pH fluctuations. (A) As we artificially increased the strength of niche overlap (by setting all off-diagonal coefficients in the interaction matrix to be a fixed negative number and making that number more negative), the sensitivity to pH fluctuations increased. This trend is disrupted when niche overlap approaches 1, because a majority of communities under such conditions lose interspecies diversity (B). As we increased the prevalence of interspecies facilitation by randomly setting a given fraction of interaction coefficients to be positive, the communities became less sensitive to pH fluctuations (C) and more diverse in species richness (D). Here pH fluctuations are sinusoidal, with *f_pH_* = 0.2/hr and ΔpH = 0.5.

## Discussion

Using empirically measured species properties, we assembled stable *in silico* communities that show coexistence of nasal bacteria. When these communities were exposed to a fluctuating pH environment, we observed that the composition of stable communities was only modestly affected. Larger pH fluctuations increased the deviation, as expected; however, even at a pH fluctuation of 1 which exceeds the observed temporal variation in the nasal passage, the composition of the majority of communities remained minimally affected. We also found that intermediate frequencies of temporal pH fluctuations caused the largest deviations in community compositions. Finally, in our results, communities with more facilitation interactions were more robust against pH fluctuations.

One important aspect of temporal fluctuations is their time scale. Even though in nature the fluctuations are not completely regular, our investigation with sinusoidal temporal fluctuations reveal the time scale at which the influence on community composition is the strongest. Our analysis reveals that the fluctuations are more impactful at intermediate frequencies between two extremes (see Methods). At very low frequencies of pH fluctuations, the community dynamics are faster than pH changes; thus we can assume the quasi-static approximation applies. In this regime, the community reaches its stable state locally (in time), and the composition follows the value of pH at any given time, regardless of the frequency of the pH fluctuations. In the other extreme, at very high frequencies of pH fluctuations, the population dynamics cannot follow rapid changes in pH and essentially the species “see” the average pH. An analysis based on the Wentzel–Kramers– Brillouin (WKB) approximation suggests that in this regime, the magnitude of change in composition (compared to the composition at the average pH) is inversely proportional to the pH fluctuation frequency. Between these two extremes is the zone that exhibits the most change in community composition with pH fluctuations (Fig 4B and Fig S7B). However, for parameters relevant to the nasal strains we are analyzing, even in this zone the changes in community composition are not drastic.

Our focus in this manuscript is on how composition of stable communities changes when environmental pH fluctuates. Another relevant question is how fluctuations in pH affect the process of community assembly. For this, we repeated the community assembly simulations (Fig S4 and S5), but under an environment in which the pH temporally fluctuated. Contrary to our expectation, the richness of resulting communities did not monotonically increase with an increase in the amplitude of pH fluctuations, regardless of fluctuation frequency (Fig S8). Instead, we found that richness only changed in a small fraction of *in silico* nasal communities. Furthermore, in cases with increased richness, *S. non-aureus* 1 (most facilitative species in our panel) was most frequently added to the community, whereas in cases with decreased richness, *S. epidermidis* (most inhibitory species in our panel) was most frequently dropped from the community (Fig S9). This observation underscores the relative importance of interaction (compared to niche partitioning) in richness outcomes in our model of nasal communities. Our finding is also consistent with predictions about augmentation and colonization resistance using a mediator-explicit model of interactions [19].

There are some limitations and simplifications in our study. First, in our investigation we have assumed that fluctuations in pH are imposed externally (e.g. by the host or the environment). It is also possible that species within the nasal community contribute to the environmental pH. Although outside the scope of this work, we speculate that if species within the community drive the pH to specific values [20,21], the impact of external temporal fluctuations of pH on community composition will be even more diminished. Second, in our model, we assumed that interactions among species remained unchanged at different environmental pH values. We examined *in silico* how pH-dependent interaction coefficients might affect our results. For this, we assumed that interaction coefficients changed linearly with pH in each case (see Methods) and asked how strong the dependency had to be to considerably change the community composition under a fluctuating pH. We observed a significant impact only when the interaction coefficients were strongly pH dependent (i.e. to the level that the sign of interactions would change within the range of pH fluctuations) (Fig S10). In the future, we plan to explore experimentally if interactions are pH-dependent and if such a dependency impacts the sensitivity of the community to temporal fluctuations in the environment.

Finally, one of the main messages of our work is that nasal microbiota is insensitive to temporal fluctuations in pH. It is tantalizing to speculate, when facing other microbial communities, under what conditions this statement is valid. Recent work by Shibasaki et al. [22] shows that under a fluctuating environment species properties play an important role in community diversity. Our results corroborate their finding. Insensitivity of the members to the environmental fluctuations— as trivial as it may sound—is a defining factor for how sensitive the community is. In the nasal microbiota, species that we are examining are adapted to the nasal environment and the range of pH fluctuations experienced in this environment is not large. As a result, the community is not majorly affected by pH fluctuations. On top of this, we also observe that interactions—in particular, facilitation and competition—can act as stabilizing or de-stabilizing factors for how the community responds to external variations. In other words, facilitation between community members acts as a composition stabilizing factor between populations, which lowers the impact of external fluctuations (Fig 6A). In contrast, inhibition between community members typically exaggerates the changes introduced by external fluctuations (Fig 6C).

## Materials and Methods

### Nasal bacterial strains

Six strains used in this study were isolated from two healthy individuals and kindly shared with us by Dr. Katherine Lemon (Table 1).

**Table 1.**
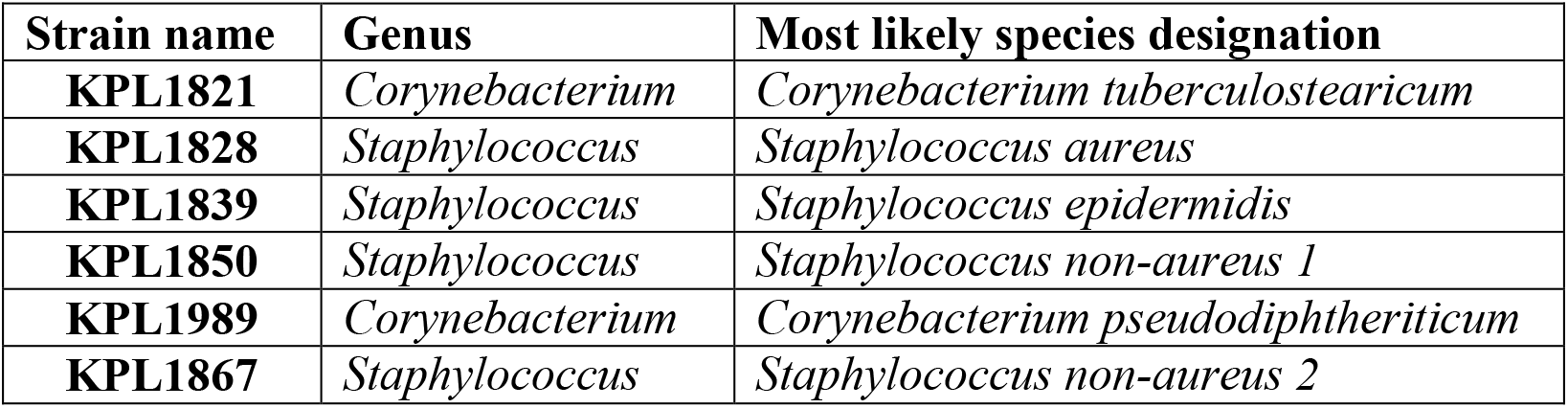
Nasal strains used in this study are listed along with their designation based on 16S rRNA gene similarity

### Cultivation conditions and medium *in vitro*

As growth medium, we have used a 10-fold dilution of the Todd-Hewitt broth with yeast extract (THY). We have diluted THY to create an environment closer to the nutrient richness of the nasal passage [23]. For collecting cell-free filtrates, cells were grown in 15 ml of media in sterile 50-ml Falcon^®^ tubes. For growth rate and carrying capacity characterizations, cells were grown in flatbottom 96-well plates. All cultures were grown at 37°C with continuous shaking.

### Characterizing the pH response of nasal isolates *in vitro*

To assess the response of nasal strains, we grew them in 10% THY after adjusting the pH within the biologically relevant range of 5.1 and 7.5 at 0.3 intervals (pH buffered with 10 g/l of MOPS). For each strain, we measured the growth rate at low population sizes (before nutrients become limiting or byproducts become inhibitory) and the final carrying capacity. These values were measured by growing replicates of each strain (typically 6 replicates) in 96-well microtiter plates incubated inside a Synergy Mx plate reader. Growth rate and carrying capacity were estimated by measuring the absorption in each well (OD_600_) at 10-minute intervals over 24 hours at 37°C. Between absorption reads, the plate was kept shaking to ensure a well-mixed environment.

### Mathematical model

To model the growth of species, we assume that in the absence of interactions, the population growth follows the logistic equations:

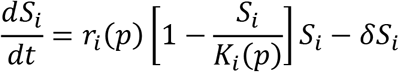

In which *r_i_*(*p*) and *K_i_*(*p*) are the pH-dependent growth rate and carrying capacity of species *i*. In our simulations, the growth rate and carrying capacity values at any given pH are found using a linear interpolation from experimentally measured values (pH 5.1 to 7.5 at 0.3 intervals). pH dependence is experimentally characterized for each strain in a monoculture, as described above, and *δ* is the dilution rate.

When multiple species are present, we assume that the presence of other species takes away resources from the environment; as a result, the growth of each species will be modulated as

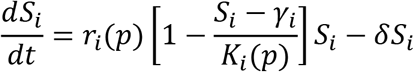

where *γ_i_* = ∑_*j≠i c_ij_*_*S_j_* and *r_i_*(*p*)*c_ij_*/*K_i_*(*p*) represents the interaction strength exerted on species *i* by species *j*. Positive values of *c_ij_* indicate growth stimulation (e.g. via facilitation by producing resources) whereas negative values of *c_ij_* indicate growth inhibition (e.g. via competition).

### Model parameters

Unless otherwise specified, the following parameters are used in the model:

| Parameter | Description | Value |
| --- | --- | --- |
| $N_c$ | Maximum number of strains for *in silico* community assembly | 20 |
| $N_s$ | Number of instances of assembly simulations run for each case | 10,000 |
| $N_{gen}$ | Number of generations simulated to obtain stable resident communities; also the number of generations simulated to assess response to environmental fluctuations | 100 |
| $pH_{rng}$ | Range of pH values (both in experiments and in simulations) | 5.1-7.5 |
| $\delta$ | Dilution rate | 0.03-0.3 hr<sup>-1</sup> |
| $N_{ext}$ | Extinction population density per species (OD) | 10<sup>-6</sup> |
| $f_p$ | Inter-strain parameter variation within each species | 20% |
| $S_0$ | Average initial cell density per strain (OD) | 10<sup>-4</sup> |
| $f_{pH}$ | Frequency of sinusoidal temporal pH fluctuations | 1 hr<sup>-1</sup> |
| $\Delta pH$ | Amplitude of sinusoidal temporal pH fluctuations | 0.5 |

### Characterizing the interspecies interactions using a supernatant assay

To characterize how species *j* affects the growth of other species *i*, we use a supernatant assay in which species *j* is grown to saturation, then all the cells are filtered out using a 0.22 μm filter (PVDF syringe filters from Thomas Scientific). The growth rate and carrying capacity of species *i* is then measured when growing in the supernatant taken from cultures of species *j*. This formulation allows us to use the experimentally measurable supernatant responses to formulate a dynamical model for mixed cultures of multiple species.

Assuming a Lotka-Volterra model, the presence of another species modulates the growth rate proportionally to the size of the interacting partner, i.e.

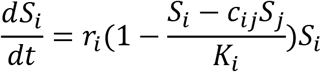

Calculating the parameters obtained from the cell-free spent media (CFSM), the carrying capacity for species *i* is reached at population *S_i,cc_* level when growth rate becomes zero, thus

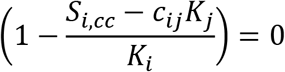

Therefore, the carrying capacity in the supernatant assay is

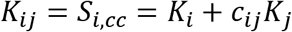

And the interaction coefficient (effect of species *j* on species *i*) can be calculated as

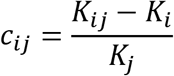

In the particular that species *i* and *j* are similar (self-effect), we have *K_ii_* = 0 and *c_ii_* = −1.

### Calculating community composition deviations

To compare community composition of a community that experienced pH fluctuation with that of the same community simulated at a fixed pH, we calculated the Bray-Curtis dissimilarity measure using the `f_dis` function (option `‘BC’`) in MATLAB^®^.

### Estimating the impact of pH fluctuations

We consider two extremes, when the fluctuations in pH are (1) much faster or (2) much slower than the population dynamics of community members. In both cases, for our formulation we define *c_ii_* = −1 and use the simplified model of populations at different pH:

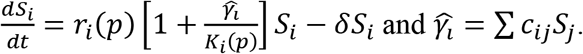

#### Case 1. Fast pH fluctuations

To estimate how the community responds under a rapidly changing pH, we use the framework of the Wentzel–Kramers–Brillouin (WKB) approximation. For the general case of 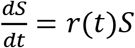, we split the population dynamics into two terms, the primary exponential term and an envelope function, *E*, for which *E*(*t*) = *e*^−*r*_0_*t*^*S*(*t*) and thus,

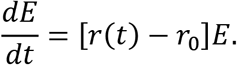

Using the WKB approximation *E* can be written using the expansion

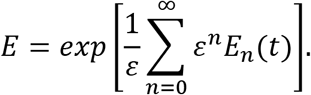

By inserting this expansion into the differential equation we obtain

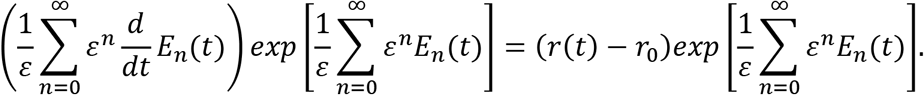

Thus

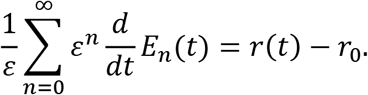

Assuming sinusoidal changes in pH, *p*(*t*) = *p_0_* + *p_d_*sin(2*πft*), to the first order, the temporal changes in growth rate can be approximated as, *r*(*t*) = *r*_0_ + *r_d_*sin(2*πft*). Therefore,

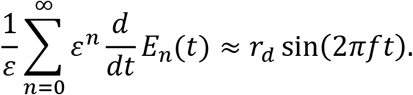

In the limit that *ε* → 0, the first terms of expansion for *E* are obtained as

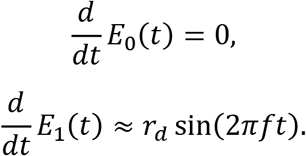

Since the continuous dilutions in our setup keeps the populations finite, *E*_0_ does not affect the solution. The dominant term for *E* thus becomes *E*_1_ and we have

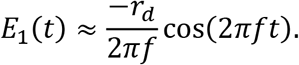

As a result,

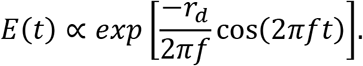

Importantly, the magnitude of change in this equation drops inversely proportional to the frequency of pH fluctuations *f*. This means that the impact of pH fluctuations diminishes at high frequencies, consistent with our intuition that in this case the community dynamics is incapable of following the environmental fluctuations and only responds to the mean value.

#### Case 2. Slow pH fluctuations

In this case, we assume the quasi-static approximation, in which fluctuations are so slow that the community approaches its steady-state at each temporal value of pH. In this situation, assuming 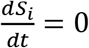, we can rearrange the equation at steady-state as

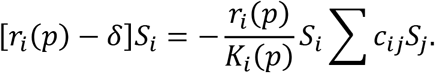

Rearranging this, we get

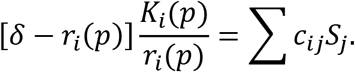

This can be written in matrix form as

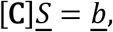

where [**C**] contains the interaction coefficients and 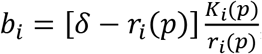; underline in our notation designates a vector. Since the interaction matrix [**C**] is pH-independent in our model, the change in composition within this quasi-static approximation can be expressed as

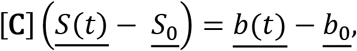

or

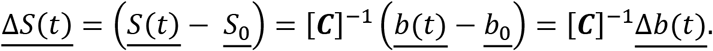

We make an additional simplifying assumption that *K_i_* and *r_i_* change similarly with pH. This leads to 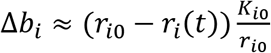. This means that the magnitude of change in community composition is the same as the change in the growth rate of species, regardless of the frequency of fluctuations, under this regime.

### Allowing pH-dependent interaction coefficients

To examine how pH-dependent interaction coefficients may affect our results, we assumed that each interaction coefficient has a linear dependence on pH with a slope (per unit pH) randomly selected from a uniform distribution in the range of [−*m*, *m*]. In other words,

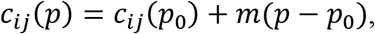

where *p*_0_=7.2 is the pH at which our characterization is performed. We examined how the community composition deviated from the reference with a fixed pH, as *m* (and thus the pH-dependency) increased.

## Acknowledgements

We would like to thank Katherine Lemon for kindly sharing the nasal bacterial strains with us. SD was supported by an NIH T32 training grant. Work in the Momeni Lab was supported by a startup fund from Boston College and by an Award for Excellence in Biomedical Research from the Smith Family Foundation.

## Code and Data Availability

Codes related to this manuscript can be found at: https://github.com/bmomeni/temporal-fluctuations.

## Supplementary Information

**Fig S1.**
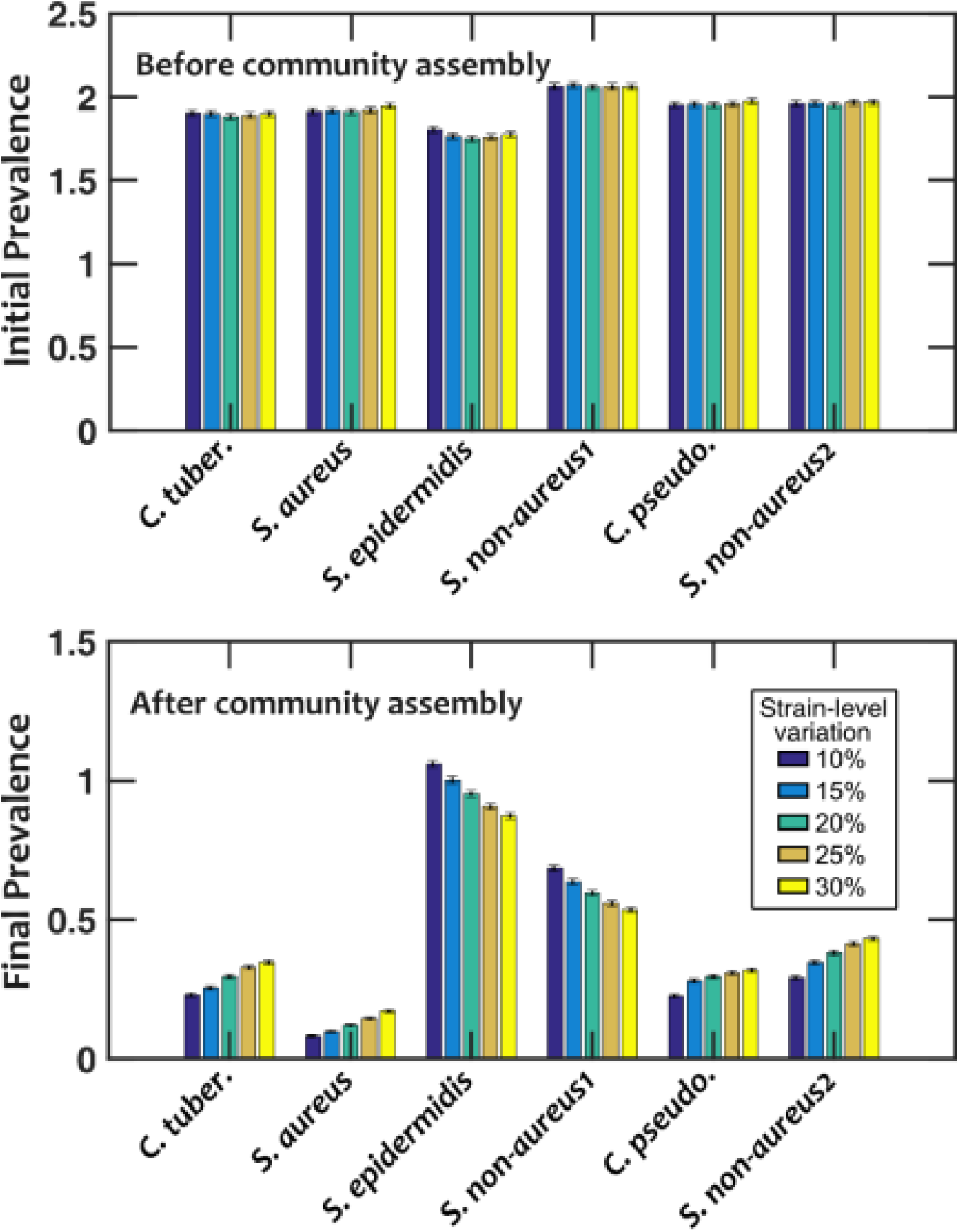
The degree of intraspecies parameter variations is chosen as a balance between strain-level and species-level diversity. Starting from communities that had a balanced representation of different strains (top), we observed that after community assembly (bottom), the prevalence of different species was affected. The prevalence here is defined as the average number of *in silico* strains from each species that appeared in each community. The pattern was more pronounced with smaller strain-level variation of 10%, and as this variation increased to 30%, the diversity between different species decreased and species prevalence became more uniform. We chose the strain-level variation of 20% to maintain some strain-level diversity, without diminishing species-level diversity. Number of *in silico* communities examined, n = 10,000.

**Fig S2.**
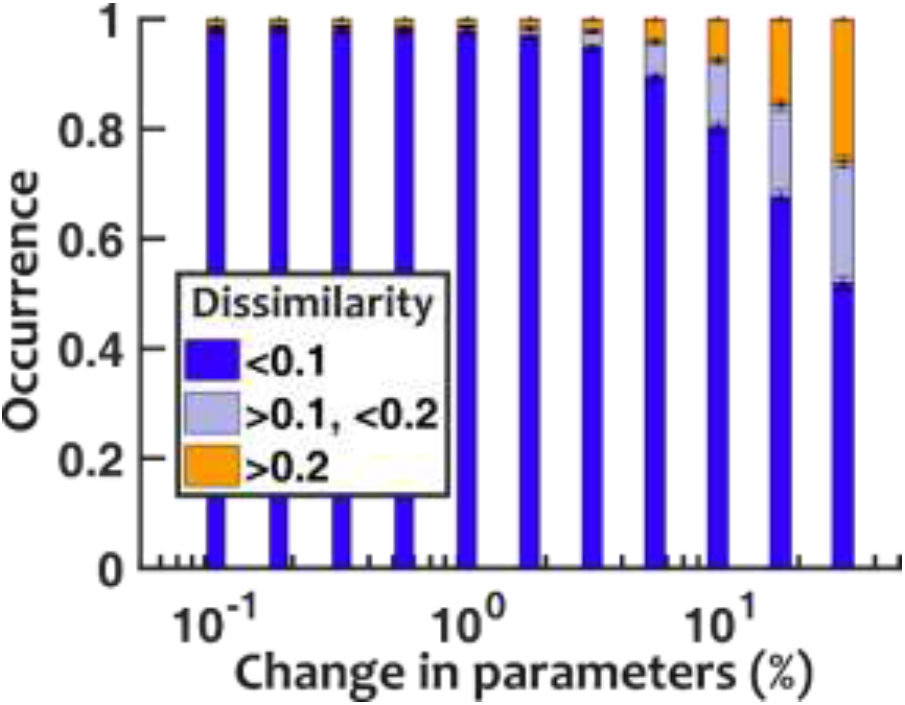
Sensitivity analysis shows that *in silico* communities are largely insensitive to measurement noise. To assess whether *in silico* communities were sensitive to experimental error in characterizing species parameters, we intentionally introduced change in measured parameters and quantified the resulting change in community composition. We observe that even when we changed all parameters by up to 10% using a uniform distribution, in silico community compositions were hardly affected. Number of cases examined, n = 10,000.

**Fig S3.**
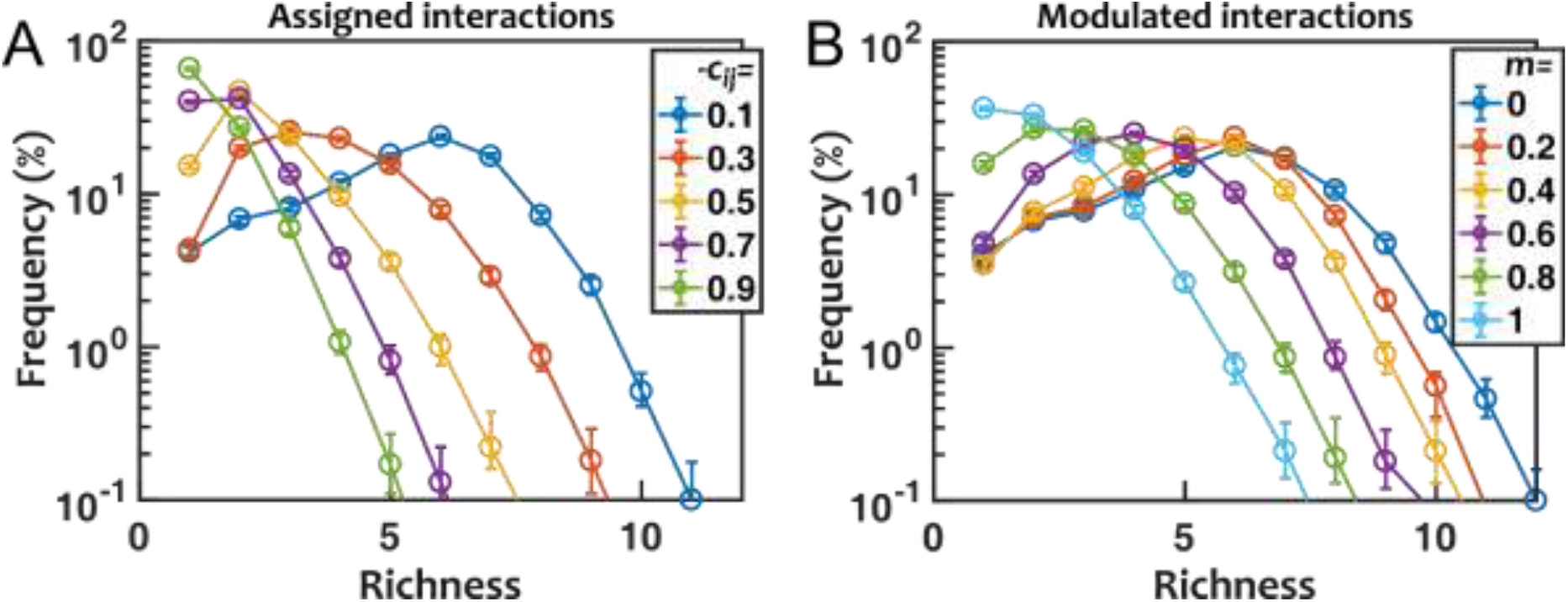
Assembly of *in silico* communities is affected by interspecies interactions. To assess the impact of interactions, in our model (A) we assigned all off-diagonal interaction coefficients to a fixed number, *c_j_*, or (B) modulated all experimentally measured off-diagonal interaction coefficients by a fixed factor, *m*. Both results show that when negative off-diagonal terms (i.e. non-self competitions) are weaker, there is more coexistence. In all cases and for all species, we assumed *c_ii_* = −1. Error-bars show bootstrap 95% confidence intervals of the mean values. Number of cases examined, n = 10,000.

**Fig S4.**
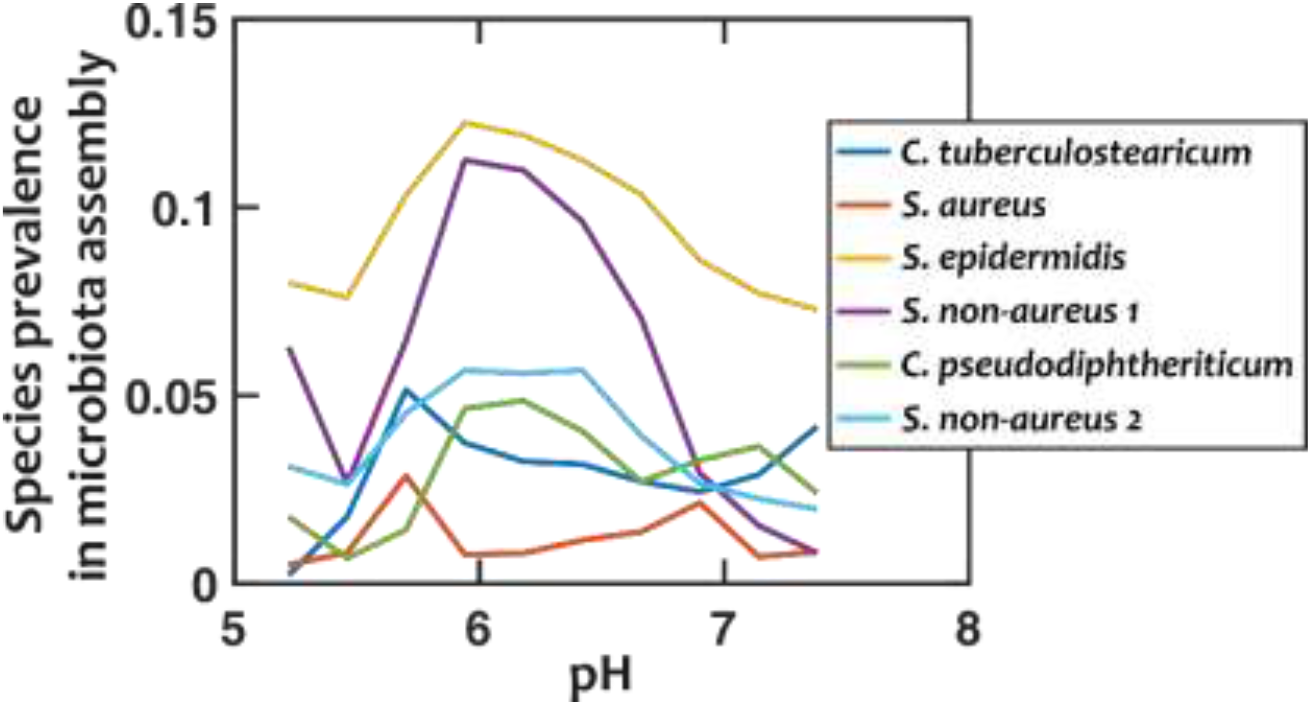
Communities assembled at different pH values show distinct and pH-dependent profiles of species-level prevalence. We binned simulated communities assembled under a fixed-pH regime according the pH and within each bin assessed how often different species appeared in assembled *in silico* communities. The prevalence of species appears to depend on pH and vary from one species to another. Number of cases examined, n = 10,000.

**Fig S5.**
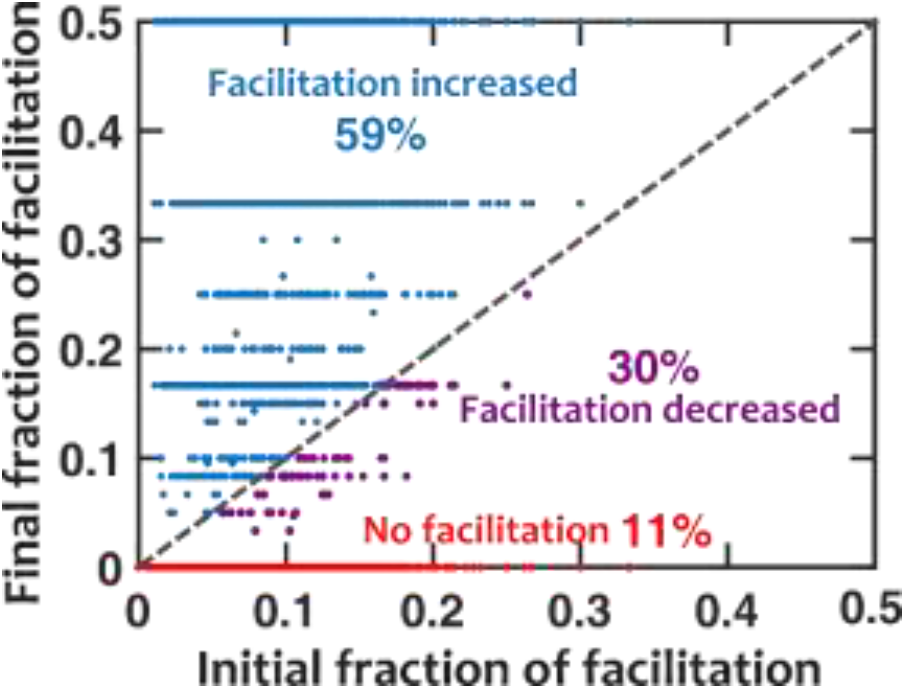
Prevalence of interspecies facilitation increases in stable *in silico* communities. Using the empirically measured parameters for species growth and interactions, we assembled communities based on *in silico* strains that resembled those species. We then examined how the fraction of interactions among final assembled in silico communities was related to the pool of species before assembly. We found that among cases that contained facilitation interactions in the initial pool of species, the final stable communities were more likely (66%:34%) to contain a higher fraction of facilitation interactions. Because there is no example of mutual facilitation among our species, the final fraction of facilitation is bound to 50%. Number of cases examined, n = 10,000.

**Fig S6.**
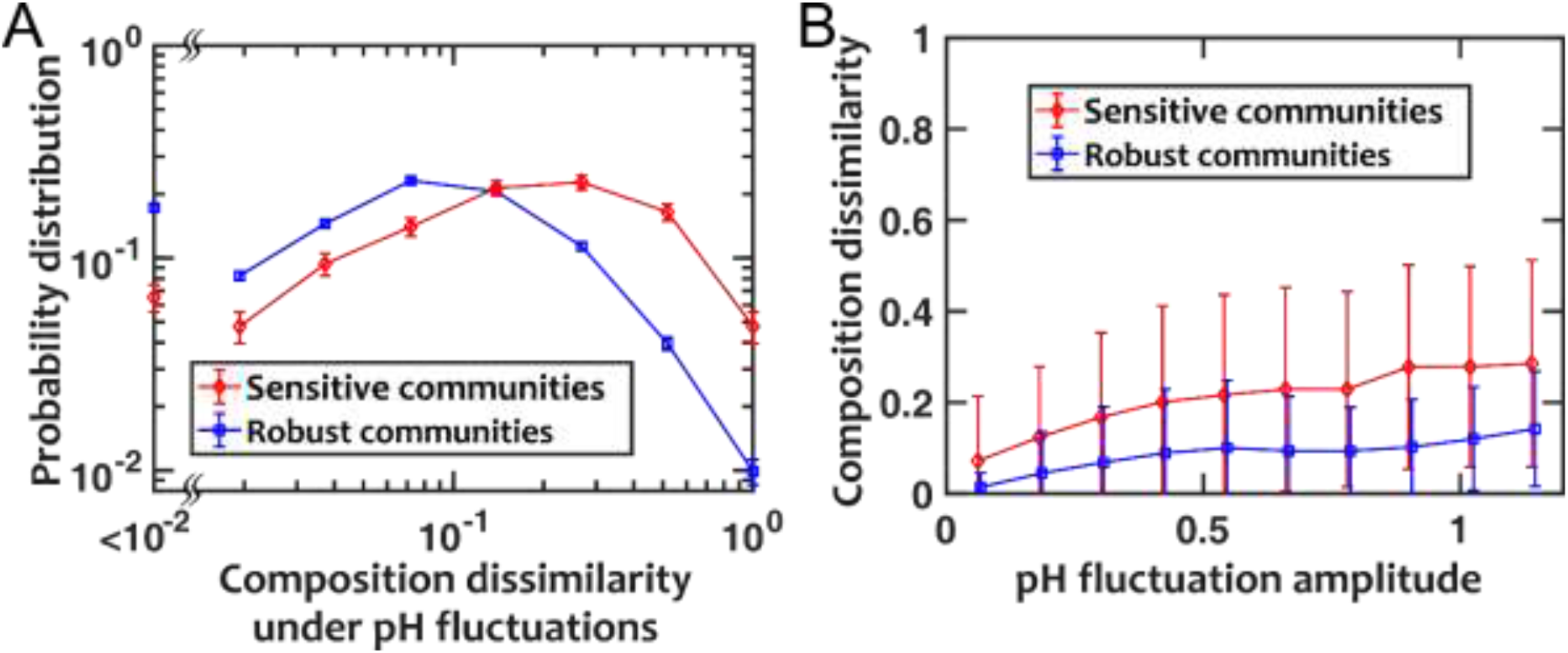
*In silico* communities that are sensitive to dilution rate changes show higher deviation when exposed to fluctuation pH compared to communities that are robust to dilution rate changes. Depending on the sensitivity of community composition to changes in dilution rate, we grouped *in silico* communities to robust (<=10% dissimilarity in composition under ±50% change in dilution rate) versus sensitive (>10% dissimilarity under ±50% change in dilution rate). When these communities were exposed to a fluctuating pH, we observed that sensitive (compared to robust) communities were more prone and showed a larger dissimilarity in composition. Among both sensitive and robust communities, dissimilarity in composition increased when the amplitude of pH fluctuations increased. Number of cases examined, n = 10,000.

**Fig S7.**
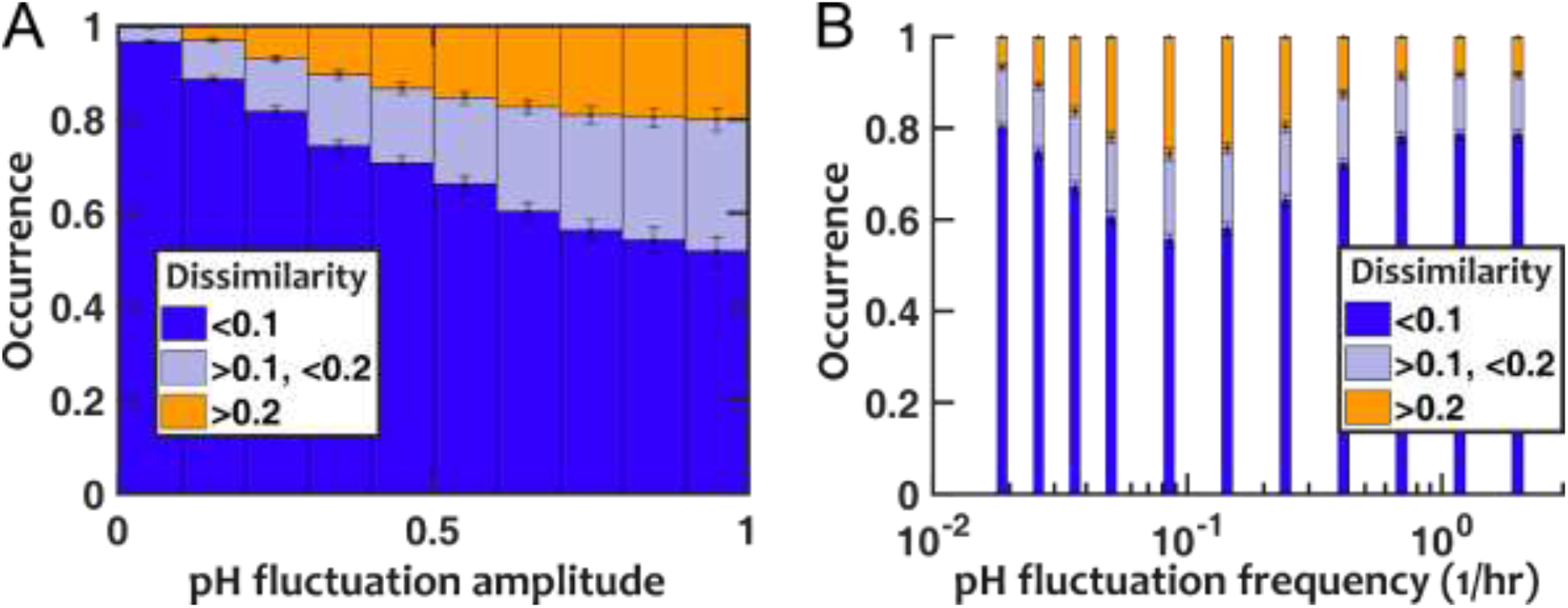
A randomly fluctuating pH shows trends similar to a sinusoidal pH variation in how composition deviations (quantified using Bray-Curtis dissimilarity) depend on the frequency and amplitude of fluctuations. Unlike sinusoidal changes in pH in the main body of the paper, here pH transitions randomly between two distinct pH values, with time between transitions following an exponential probability distribution (i.e. transition times following a Poisson distribution). In these results, the average frequency of pH transitions is *f_pH_* = 0.2/hr and the step between the two pH levels is ΔpH = 0.5 (on average comparable to ΔpH = 0.5 for sinusoidal fluctuations).

**Fig S8.**
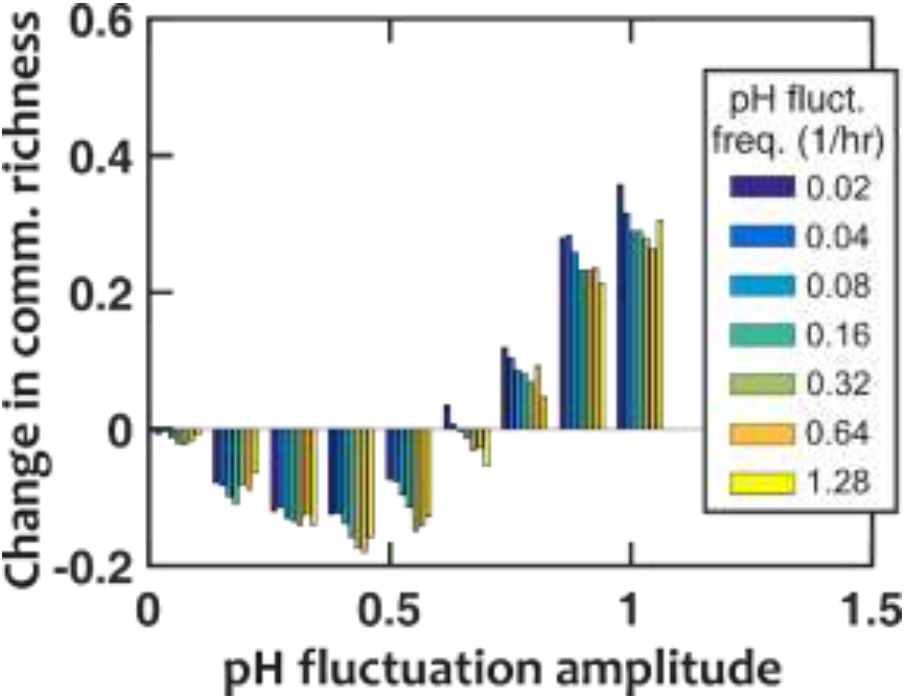
If pH fluctuates during community assembly, the richness of resulting communities could be different from the fixed-pH case. However, unlike theoretical predictions, the richness does not monotonically increase when the amplitude of pH fluctuations increases. We examined fluctuations at different frequencies and observed little dependence of the outcome on the pH fluctuation frequencies. Nasal microbiota assembly is only modestly affected under pH fluctuations. Number of assembly cases at each fluctuation frequency, n = 1,000.

**Fig S9.**
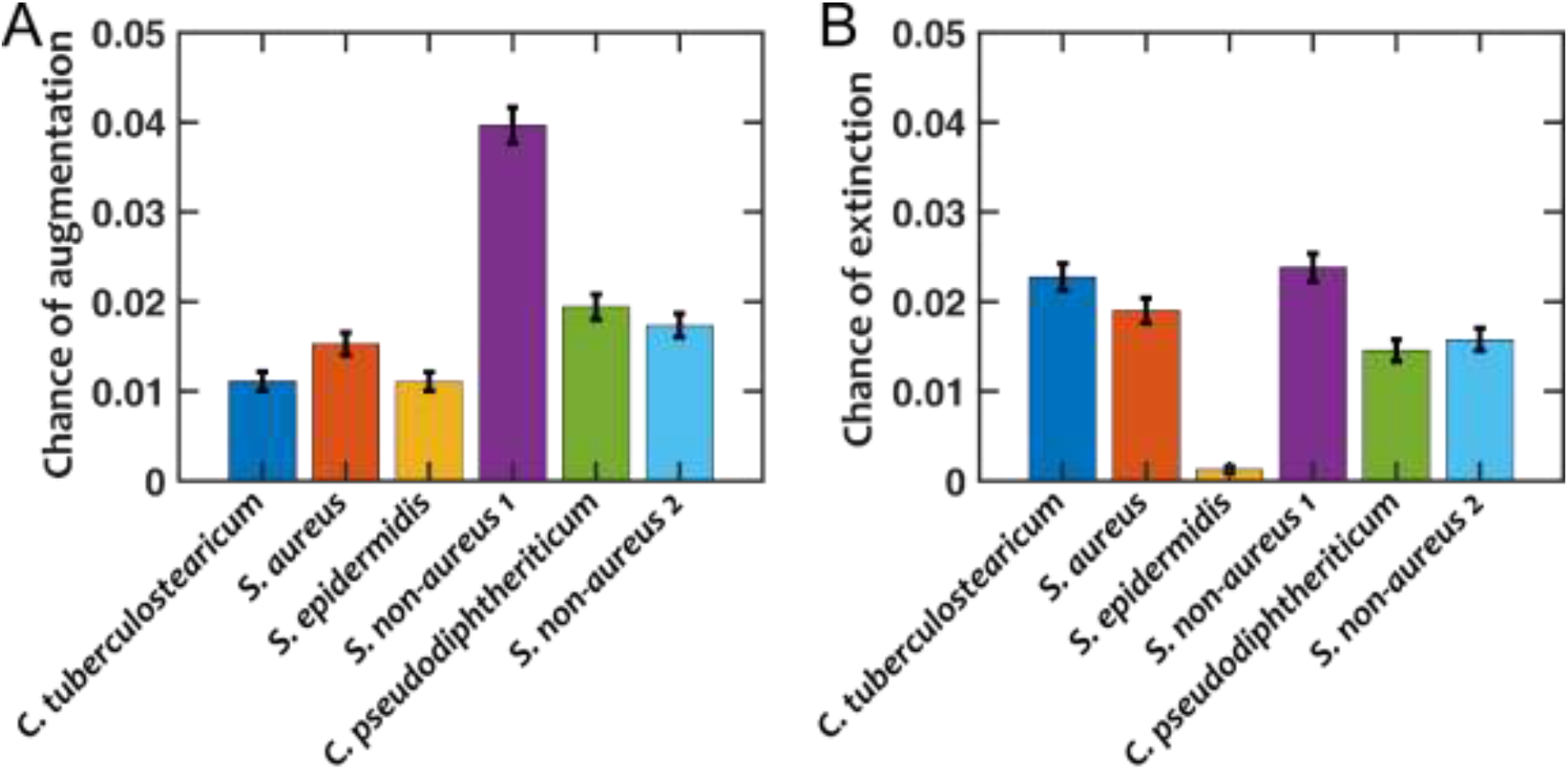
Species with the most facilitation are most likely to be added into and species with the most inhibition are least likely to drop from nasal microbiota that are assembled under a fluctuating pH. We enumerated the probability at the species level that a member would be (A) added to or (B) dropped from a community during assembly when the pH fluctuated (compared to the fixed-pH case). The amplitude of pH fluctuations tested varied from 0.1 to 1 and the frequency of fluctuations tested varied from 0.02/hr to 1.28/hr (similar to Fig S8). Number of assembly cases at each fluctuation frequency, n = 1,000.

**Fig S10.**
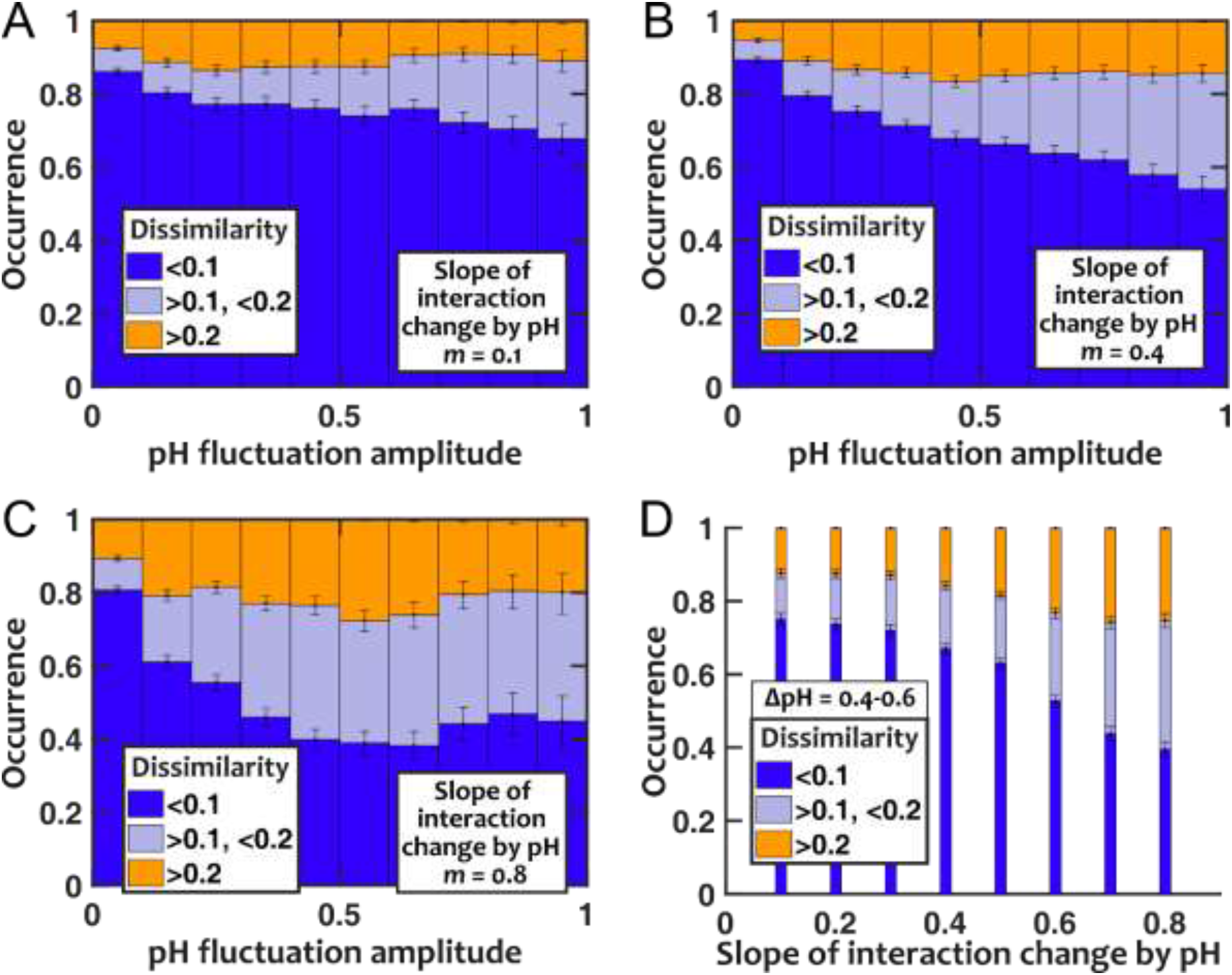
With temporal fluctuations in pH, pH-dependency of interaction coefficients affect the community composition only when the dependency is very strong. We allowed interaction coefficients to each linearly change with pH with a slope randomly selected from a uniform distribution in the range [−*m*, *m*]. (A-C) Under a temporally fluctuating pH, as the strength of pH dependency of interactions (i.e. *m*) is increased, we observe more deviation from the stable community composition. For each *m* in (A-C), the composition deviates more as the amplitude of pH fluctuation increases, consistent with our results for pH-independent interactions in Fig 4. (D) Comparing the results for simulated cases with a pH fluctuation amplitude (ΔpH) between 0.4 and 0.6, we note that the effect of pH-dependent interactions on composition deviation is minimal until the dependency becomes strong enough that the change becomes comparable to the interaction coefficients themselves (*m* > 0.4). The frequency of fluctuations *f_pH_* = 0.2/hr (similar to Fig 4). Number of communities examined, n = 10,000.

